# Mutational Analysis of an Intrinsically Disordered Region in the *E. coli* Phosphatase CheZ, a Regulator of Chemotaxis

**DOI:** 10.1101/607010

**Authors:** Amber Ward, Marilyne Kamegne, Paulette Dillard, John T. Moore

## Abstract

As a model for Intrinsically Disordered Region (IDR) structure-function, we investigated the IDR present in the *E. coli* chemotaxis regulatory phosphatase CheZ. The CheZ IDR (amino acids 169-200) is attractive for study because CheZ is part of an exceptionally well characterized and easily tractable system and has an available high-resolution crystal structure. The CheZ IDR contains striking evolutionarily conserved regions that are functionally critical as shown by the fact that changes in single specific amino acids in the conserved IDR regions are able to abolish normal chemotaxis. We have focused on identifying suppressor mutations in the coding region of CheZ G188E, a variant with a >80% reduction in chemotactic swarm activity. Our screen identified 6 suppressor mutations that restored swarm activity to wild-type levels. Interestingly, the suppressor mutations were found not in the CheZ coding region, but in CheY, the phosphoprotein substrate of the CheZ phosphatase. The 6 suppressor mutations were restricted to two amino acid codons. Three suppressor mutations were found at position CheY A42 and three at position CheY M60. A model for how the changes in CheY restore CheZ activity is presented. These studies in dissection of the CheZ IDR will be useful in extending our understanding of IDR structure-function. The accumulating knowledge of IDR principles will be a significant component of a broadening understanding of protein biophysics with direct implications in basic protein structure and protein engineering research.

## Introduction

### IDRs and protein function

Traditionally, protein function has been viewed as dependent on the folded three-dimensional structure of a polypeptide. This stems from the fact that most of our knowledge of protein function comes from the study of stable, compact ordered structures that can be visualized using crystallography. Recently, however, as available protein sequences expand through whole genome analysis and methods to predict protein disorder have been developed (Dunker *et al*., 2015), we are now discovering that, in addition to folded domains, a large percentage of the proteome is comprised of intrinsically disordered regions (or IDRs, protein sequences that lack a fixed or ordered three-dimensional structure). It has been estimated that 44% of human protein-coding genes encode disordered segments of >30 amino acids in length or longer (He *et al*., 2009).

Proteins can be characterized along a spectrum, from completely structured to completely disordered (van der Lee *et al*., 2014a). Most proteins lie in the middle, with a modular architecture containing IDRs interspersed with structured regions. Both the structured and disordered regions are important to achieve the diverse range of protein functions (Dunker *et al*., 2015; He *et al*., 2009). With the recognition of the prevalence and importance of IDRs, there has been a recent surge of interest in their biochemistry and function (for review, see Oldfield and Dunker, 2014).

Because of their dynamic nature, IDRs can be active participants in protein function (Gokhale and Khosala, 2000; Ma *et al*., 2011). IDRs have been shown to be involved in diverse functions including cell-cell recognition, cellular signaling pathways, protein partner recognition, and protein half-life determination (Borcherds *et al*., 2013, van der Lee *et al*., 2014b). The conformational disorder of these domains can impact *flexibility* (allow movement of linkers and of domains relative to each other), and *spacing* (regulate the distance between domains). These properties have recently been shown to be important in the function of the chemotaxis kinase CheA, a protein which contains two IDRs (Wang *et al*, 2012; Wang *et al*, 2014).

Because of the high degree of characterization of *E. coli* chemotaxis and the proteins involved, as well as the ease of which chemotaxis function is assessed in phenotypic assays, we have focused on proteins involved in chemotaxis to study IDRs.

### Chemotaxis

In *Escherichia coli*, chemotaxis is governed by regulating a shift between two distinct swimming patterns, one generated by counterclockwise (CCW) rotation of the flagella and one generated by clockwise (CW) (for review, see Parkinson, 1993; Amsler and Matsumura, 1995; Bi and Sourjik, 2018). CCW rotation results in the *smooth swimming* of the cell in one direction, with the flagella forming a stable flagellar bundle behind the cells. When the flagella switch to CW rotation, the bundle is no longer stable and the flagella separate and forward motion ceases. Normal chemotaxis behavior depends on a critical balance between the CCW and CW states. The switch between CW rotation (tumble) and CCW (smooth swimming) in response to changes in the chemical environment is controlled by the chemotaxis signal transduction pathway.

### CheY and CheZ are key regulatory proteins in the chemotaxis signal transduction pathway

The presence of changing concentrations of a chemo-attractant or chemo-repellant bound to transmembrane chemoreceptors modulates the rate of autophosphorylation of the chemotaxis CheA kinase (Baker *et al*., 2006). A phosphoryl group is transferred from CheA-P to the response regulator CheY to form CheY-P. CheY-P binds to the base of the flagellum and stabilizes the CW flagellar state. The intracellular concentration of CheY-P dictates the rotational state of the flagella. Another interacting partner, CheZ, accelerates the dephosphorylation of CheY-P 100x over its intrinsic autodephosphorylation rate. Thus, the intra-cellular concentrations of CheY-P are regulated by a balance of incoming phosphoryl groups from the kinase CheA and outgoing phosphoryl groups resulting from CheY autodephosphorylation and dephosphorylation mediated by CheZ.

Our studies will focus on an IDR in the chemotaxis protein CheZ. The crystal structure of CheZ complexed with CheY bond to the stable phosphoryl group analog 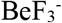 (Zhao *et al*., 2002) was used to guide these studies. CheZ is a helical homodimer (Figure 1, dark and light blue ribbons) that binds two 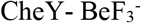 molecules (green ribbons). The CheZ dimer is comprised of two types of structured domains: (i) The N-terminal domain (CheZ_N_) is dominated by a long four-helix bundle comprised of a helical hairpin from each CheZ chain and (ii) the C-peptide (CheZ_C_) is an alpha helix comprised of the C-terminal 15 residues of each CheZ chain. Each CheY molecule interacts with two independent surfaces on CheZ: one binding surface is the entire CheZ_C_ helix and the other binding surface is a region on CheZ_N_. The structural domains containing the two CheZ binding surfaces for CheY (CheZ_N_ and CheZ_C_) are connected by the 32-residue CheZ IDR, which was not visible in the crystal structure. Catalysis (dephosphorylation of CheY) occurs as a result of the interaction of CheY-P with CheZ_N_.

**Figure 1:**
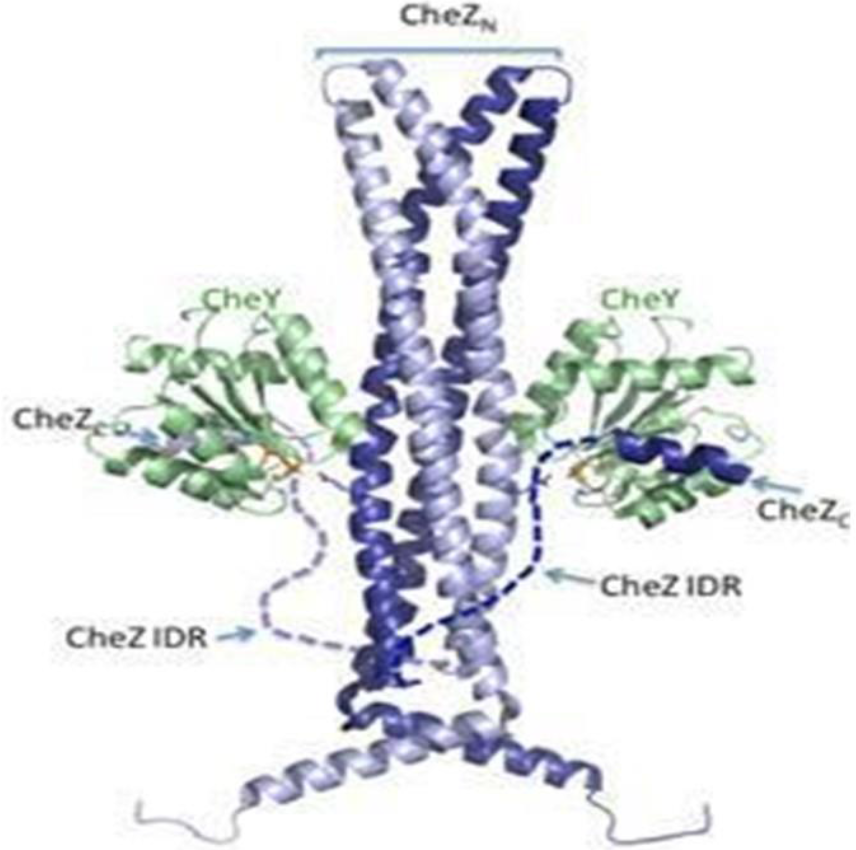
Crystal structure of CheZ: Activated Chey heterodimer (Zhao et al., 2002). Chez is a helical homodimer (Figure 1, dark and light blue ribbons) that binds two Chey molecules (green ribbons). The position of the IDR in the Chez is indicated. This structure is used with permission from the lab of Dr. Robert Bourret, UNC-Chapel Hill. 10.2210/pdb1KMl/pdb

There are several observations that serve as evidence that the CheZ IDR carries out critical function in CheZ activity. Of the two CheZ binding surfaces for CheY-P, CheZ_C_ provides the bulk of the binding energy (Blat and Eisenbach, 1996; Guhaniyogi *et al*., 2006, Guhaniyogi *et al*., 2008). However, there is no CheZ-mediated dephosphorylation of CheY-P when CheZ_C_ and CheZ_N_ are present as separate proteins (Zhao, 2002). Additionally, there are single site IDR mutants that greatly reduce CheZ activity (Boesch *et al*., 2000). For example, a G188E mutation in the IDR has been reported to significantly reduce activity (Blat and Eisenbach, 1996; Boesch *et al*., 2000).

A model of association of CheY-P with CheZ involves initial binding of CheY-P to the CheZ_C_ helix. The CheY-P is then “reeled in” to allow interaction with CheZ_N_ so that catalysis can occur (see schematic representation in Figure 2, and Silversmith, 2005; Freeman *et al*., 2011). This “reeling in” involves localization of the bound CheY-P in the vicinity of the binding surface on CheZ_N_. The positive cooperativity displayed by CheZ with respect to CheY-P and other kinetic data (Silversmith *et al*., 2008) indicate that association of the second CheY-P of the dimer is faster than association of the first. It is currently unknown whether the IDR plays a role in this process, but it is conceivable that CheY-P association rate could be modulated by IDR dynamics that affect the accessibility of CheZ_C_ and/or CheZ_N_. However, since little is known about CheZ structure in the absence of CheY and the CheZ IDR is not visible in the crystallographic images, important functional characteristics of the sequence, such as bends or kinks, relatively rigid structures, bending or wrapping around other domains, or direct interactions with other regions of CheY/CheZ are unknown.

**Figure 2:**
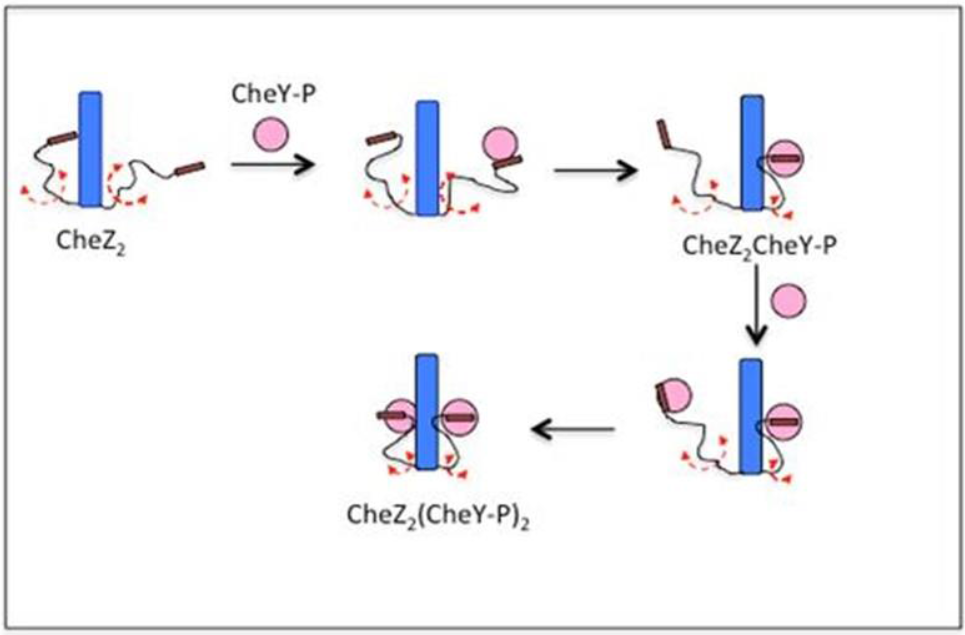
Schematic representation of a model for binding of activated Chey (CheY-P, pink circles) to Chez (blue bars) and its facilitation by the Chez IDR (black lines). According to this model, association of CheY-P with Chez involves initial binding of CheY-P to the CheZ_C_ helix. The CheY-P is then “reeled in” to allow interaction with CheZ_N_ so that catalysis can occur. This “reeling in” involves localization of the bound CheY-P in the vicinity of the binding surface on CheZ_N_ and in this model, this process is accomplished by the IDR. Figure provided by R. Silversmith (University of North Carolina, Chapel Hill) and used with permission.

We plan to approach the question of the specific role played by the CheZ IDR in binding activated CheY through genetic experiments. The goal of these studies is to define the basic functional constraints and characteristics of the CheZ IDR as well as to identify sequences in both CheZ and CheY that contribute to IDR function. This work will set the stage for additional genetic and biophysical analyses to follow.

## Materials and Methods

### Strains and plasmids used in these studies

*E. coli* strains were obtained from the lab of Drs. Robert Bourret AND Ruth Silversmith (University of North Carolina Chapel Hill, Department of Microbiology and Immunology). *E. coli* strains used in chemotaxis studies are described in Boesch *et al*., 2000, and Freeman *et al*., 2011. *E. coli* RP5231 (∆*cheYZ* strain) was transformed with plasmid pRS3 which contains an operon expressing both CheZ and CheY. The “wild-type” CheZ on pRS3 contains an inadvertent E134K mutation which reduces activity in chemotaxis swarm assay by 20% (Boesch *et al*., 2000). The mutator strain used in suppressor screen (NR9458) is described in Schaaper and Cornacchio, 1992.

### Swarm Assay

The swarm assay (Wolfe and Berg, 1989) was performed essentially as previously described (Boesch *et al*., 2000; Freeman *et al*., 2011). To measure chemotaxis activity, bacteria from plated colonies were spotted onto a swarm agar plate (1% [wt/vol] tryptone, 0.5% [wt/vol] NaCl, 0.3% [wt/vol] BactoAgar) using a sterile toothpick. Chemotactic swarming requires a balance between CW (high CheY-P) and CCW (low CheY-P) behavior. Thus a shift of CheZ activity downwards (loss-of-function) or upwards (gain-of-function) both will abolish normal activity. In this assay, a seeded bacterial population spreads away from the initial seeded location across an agar surface, and the rate (swarm diameter versus time) is dependent on nutrient composition and viscosity of culture medium. Each swarm plate contained a wild-type strain positive control and a negative control (an *E. coli* strain deleted for the *cheY*/*cheZ* genes). Growth of the control and test bacteria was allowed to proceed for 12 hours at 30 °C. Plates were subsequently photographed and diameter of colonies measured (swarm diameter). The chemotaxis rate was expressed as percent of wild type (diameter of mutant/diameter of wild type). The output from these studies provides a semi-quantitative rank-order of relative chemotaxis phenotypes.

### Tethered Cell Assay

The tethering assay has been described (Bearinger *et al*., 2009). Cells derived from individual colonies were grown overnight at 30 °C in tryptone broth to exponential growth phase, harvested, and then passed through hypodermic needle to shear flagella bundles from their base. Treated bacteria were then affixed to anti-flagellar antibody (gift of Dr. Robert Bourret, University of North Carolina) on a cover slip in a buffer supplied with lactate as an energy source followed by microscopic visualization of rotating bacteria. Ab-stabilized bacteria were visualized at 400x magnification on an Olympus microscope and imaged using 2-minute video clips (Axio digital camera). Mutant strains were characterized with respect to the degree of CW versus CCW flagellar rotation (from + to ++++, ++++ meaning exclusive CW rotation).

### Suppressor Screen

The CheZ mutant suppressor screen was carried out using an established procedure (Boesch *et al*., 2000; Freeman *et al*., 2011). Plasmid pRS3 (carrying wild type *cheYZ*) was transformed and grown in *E. coli* NR9458 which carries the *mutD5* allele (exhibits mutation frequencies 50-100 times higher than wild-type when the bacteria are grown in minimal media). Transformation and culture times were carried out as described previously (Boesch *et al*., 2000). Transformation cultures of the mutagenesis NR9458 strain were plated on selective agar plates and plasmids were isolated and subsequently transformed into the motility strain *E. coli* RP5231 (∆*cheYZ* strain). An aliquot of the RP5231 transformation mix was inoculated into a larger volume and amplified overnight.

To screen for suppressors, an aliquot of the 100ul overnight culture was streaked across the center of swarm plates and incubated for 22h at 30°. Transformants were screened for normal or improved chemotactic behavior relative to negative controls (the majority of bacteria in the streaked regions of the plate) on the same plate. Potential suppressor mutants were picked from the periphery of the swarm rings, single colony purified, and DNA purified and re-transformed into fresh swarm host strain. The resultant clones were then re-assessed in swarm and tethered cell assays. Plasmid DNA was purified from colonies of interest for sequence analysis.

### Sequence Analysis

Purified miniprep plasmid DNA (Qiagen Corporation) and accompanying *CheY/CheZ* sequencing oligos were submitted to GeneWiz (Morrrisville, NC) for Sanger sequence analysis.

## Results

### 1. Identification of highly conserved amino acids in the CheZ IDR

A sequence alignment of > 200 CheZ proteins across a non-redundant protein dataset (pfam analysis) was carried out. The results for the IDR region are shown in Figure 3. An indication that the CheZ IDR plays a more complex role in CheZ function beyond merely providing a connection between the 4-helix bundle and the C-helix is the presence of certain amino acids, in particular a Gly-Pro pairing (amino acids 188-189 in *E. coli* CheZ), that displays strong evolutionary conservation. The Gly-Pro motif (aa188-189) in the CheZ IDR is strictly conserved across species with a CheY: CheZ pathway analogous to the *E. coli* pathway. Other conserved residues were also noted such as a conserved Val-Val motif at the carboxyl-terminal end of the CheZ IDR. To the best of our knowledge, a specific function for any of these conserved regions has not been described previously in the literature.

**Figure 3:**
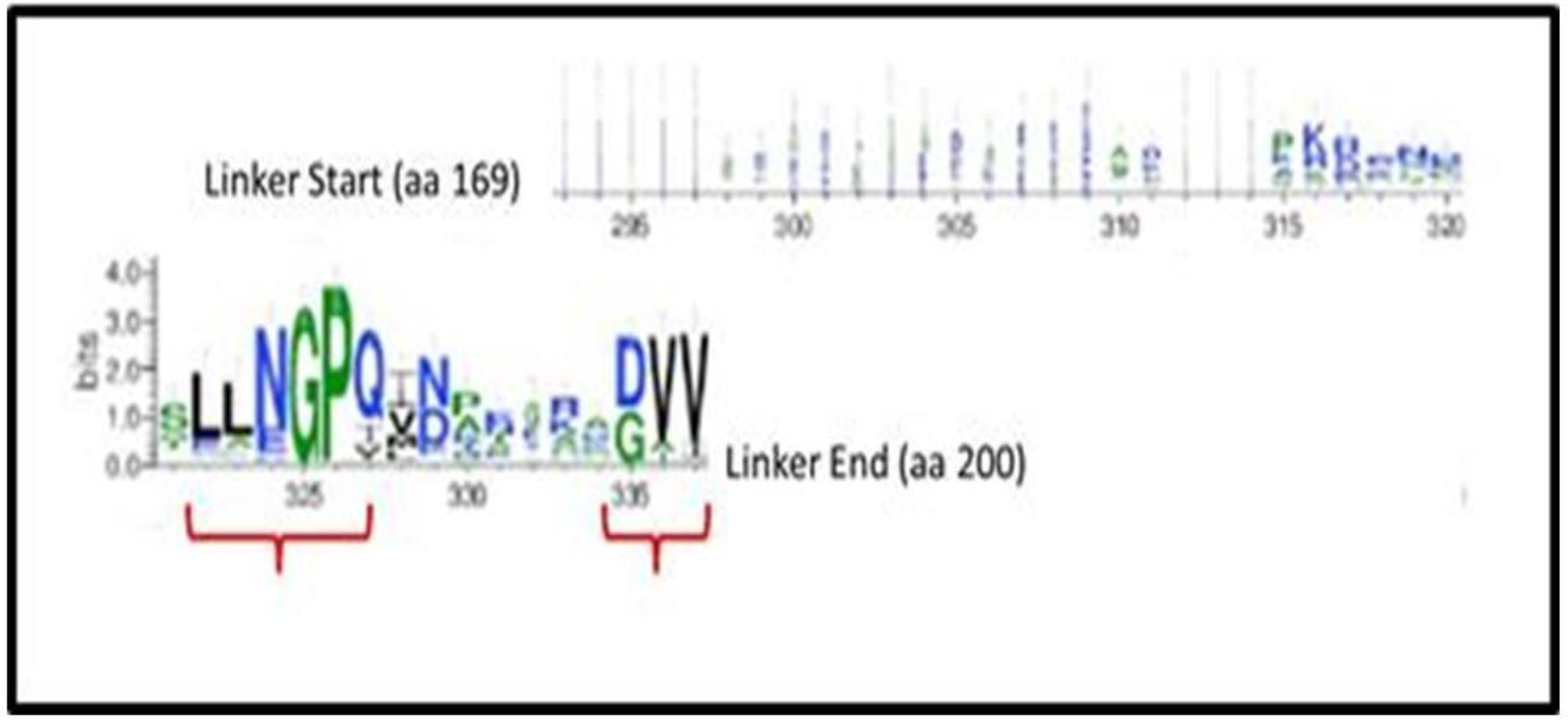
A sequence comparison of Chez proteins across a non-redundant protein dataset (pfam analysis) was carried out and highlights that the Gly-Pro motif (aa188-189) is the most conserved sequence in the in the Chez IDR (relative size of letter indicates relative degree of conservation). The Gly-Pro motif is strictly conserved across species with CheY:CheZ pathway analogous to the *E. coli*. Other conserved residues were also noted such as a conserved Val-Val motif at the carboxyl-terminal end if the Chez IDR. Figure provided by R. Silversmith (University of North Carolina, Chapel Hill) and used with permission.

### 2. Assessment of phenotype of CheZ linker mutation (G188E) in chemotaxis assays

#### Swarm assay

The CheZ G188E mutant was isolated in a prior genetic screen and their phenotype assessed (Boesch *et al*., 2000). The swarm assay was repeated in our studies and the results are completely consistent with the above showing no detectable swarm activity compared to a negative control after 9h (not shown) and ≤10% wild type after overnight incubation at 30C (Figure 4). Swarm activity detected in P189L was slightly higher than that seen in G188E, but significantly reduced versus the wild-type control (Figure 4).

**Figure 4:**
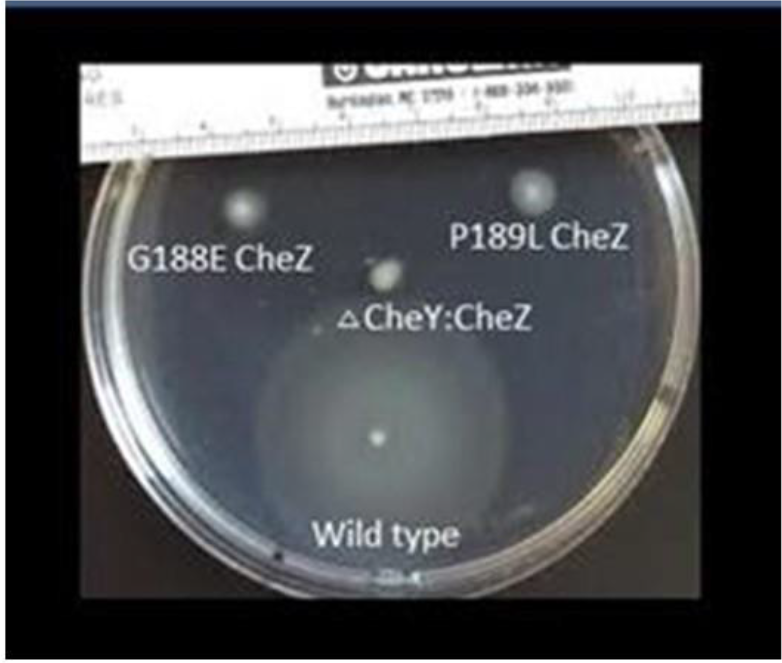
Swarm assay comparing four E coli strains: 1) wild-type (positive control), 2) strain with deleted CheY;CheZ genes (delta CheY:CheZ, negative control), 3) strain expressing Chez G188E, and 4) strain expressing Chez P189L.

#### Tethered cell assay

The tethered cell assay was used to provide a direct measure of the relative clock-wise (CW) versus counter-clockwise (CCW) rotation of the flagella motor. Because CheY-P induces CW rotation, the % of time spent in CW rotation is a surrogate marker of relative CheY-P levels. The tethered cells assay can be used to assess whether chemotaxis defects detected in the swarm assay are due to increased CheZ activity (gain-of-function) or decreased CheZ activity (loss-of-function). Chemotaxis depends on a balance between tumbling and smooth swimming behavior, so either gain-of-function mutations, resulting in increased CCW flagellar rotation and smooth swimming behavior, or loss-of-function mutations resulting in increased CW flagellar rotation and increased tumbling behavior, could result in loss of chemotaxis.

In this assay, the CheZ G188E mutation was scored as ++++ (exclusively CW rotation) and the P189L mutation as +++ (predominantly CW rotation), thus indicating both mutations represent loss-of-function mutations in CheZ. These data are consistent with the tethered cell assay data generated by Boesch *et al*., 2000. Boesch *et al*. 2000 reported that neither displayed detectable swarm diameter after 9h at 30C and either exclusive CW rotation in the tethered cell assay (G188E) or majority CW rotation with infrequent reversal to CCW (P189L). These data are also consistent data from Wolf and Berg (1989) that show that predominant CCW mutants migrate less in agar than CW mutants.

### 3. Identification of suppressors of mutants in the CheZ IDR coding region

We carried out a suppressor screen to identify amino acid changes in the CheY; CheZ complex that would restore the defective swarm assay phenotype of CheZ G188E. Candidate suppressor clones exhibiting increased swarm activity relative to the mass of other cells in the screen were picked (Figure 5, panel A). After re-streaking the selected clones to get single isolated colonies, plasmid DNA was isolated from these bacteria and re-transformed into a fresh host cell line and clones were then re-assessed for swarm activity (Figure 5, Panel B). Swarm diameters were assessed at a single time point (12h) and thus data on swarm activity is not considered quantitative. Colonies with swarm activity restored to near wild-type levels in this assay were chosen for further study. Note that suppressor colonies exhibited the concentric circles characteristic of normal chemotactic swarming behavior (Figure 5, panel B), indicating that they did not represent a “pseudotaxis” phenotype, defined as migration of nonchemotactic cells in porous agar (Wolf and Berg, 1989). Plasmid DNA from the confirmed clones with higher swarm activity was purified and submitted for sequencing. Results from sequencing showed that the clones with increased activity were a combination of revertants (22 clones) and suppressors (6 clones).

**Figure 5:**
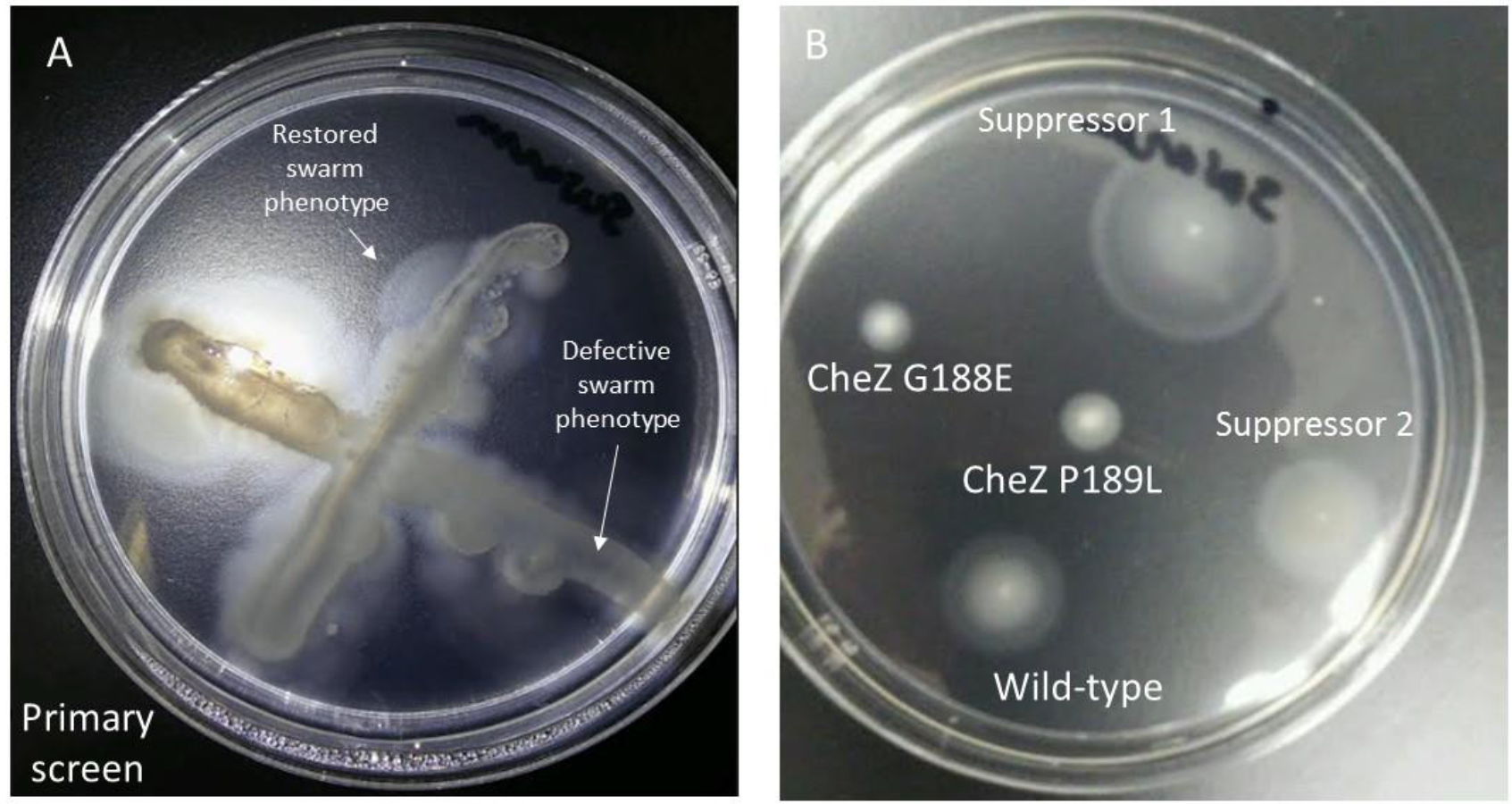
Suppressor screen and isolated suppressor colonies were compared on swarm plates. Panel A. An example of one of the primary screen plates is shown. Colonies that extended outward from the mass of bacteria perpendicularly streaked across the middle were picked from their outer edge and re-streaked (multiple times) to isolate single colonies. Plasmid DNA from the single colonies was used to re-transform new host bacteria. Panel B. Single suppressor clones were tested in the swarm assay along with relevant positive and negative controls.

#### Revertants

As described in the Materials and Methods section, the original “wild-type” plasmid used in the studies was a variant of CheZ containing an inadvertent E134K mutation. This variant has been shown previously to have approximately 20% less activity in the swarm assay versus E134 (Boesch *et al*., 2002). Interesting, of the 22 revertants we identified in our screen, 16 of the 22 were double revertants (reversion of both G188E and E134K). This result indicates that this suppressor screening method is a highly efficient method for identifying subtle improvements in chemotactic behavior, since we were able to strongly bias for clones with only 20% more swarm activity that the expected majority of revertants (single reversion of E188 back to G). In our primary screen, we intentionally selected colonies with the largest diameters, thus biasing toward stronger phenotypes. This methodology may be useful in screens seeking to fully optimize a phenotype or identify small, incremental increases in chemotaxis pathway protein activity.

#### Suppressors

A total of 6 suppressor mutants were identified in our screen and all were contained in CheY coding region (Table 1). Three of the suppressor mutations were localized to codon CheY A42 and three in CheY M60. The nucleotide changes resulted in two amino acid substitutions at A42 (A42V and A42T) and two at M60 (M60L and M60V). The existence of two amino acids at each position indicates at least two independent mutations occurred at each codon to generate the suppressors. All the suppressor mutations resulted from single nucleotide changes relative to original G188E clone. With the exception of the polar Thr residue at CheY A42, the new amino acids were hydrophobic residues, as expected from the amino acid substitutions possible via a single nucleotide changes.

**Table 1:** CheY suppressor clone sequences: Suppressor clone amino acid mutations and their associated nucleotide changes are shown in red. Unchanged amino acid and nucleotide sequences are shown in black.

| Clone | AA | Nucleotide | AA | Nucleotide |
| --- | --- | --- | --- | --- |
| 1 | A42 | GCT | **M60L** | **TTG** |
| 2 | A42 | GCT | **M60L** | **TTG** |
| 3 | A42 | GCT | **M60V** | **GTG** |
| 4 | **A42V** | **GTT** | M60 | ATG |
| 5 | **A42V** | **GTT** | M60 | ATG |
| 6 | **A42T** | **ACT** | M60 | ATG |

## Discussion

IDRs play important functional roles in proteins but are poorly understood. Since IDRs are typically not visible in the most common tool used to study protein structure (crystallography), non-crystallographic methods are needed to provide insight into key regions of IDRs and their specific functions. We have used a genetic method (mutation analysis and accompanying suppressor screening) to investigate a highly conserved amino region of the CheZ IDR.

The strongly conserved Gly-Pro motif (aa188-189) in the CheZ IDR provided an initial focus for these genetic studies. Our suppressor screen of CheZ G188E identified mutations which restored activity, but interestingly none of the suppressor mutations were located in the CheZ coding region, but in CheY instead. The fact that the suppressor mutations were found in CheY and not CheZ is intriguing, but not totally unanticipated. CheY and CheZ are closely interacting partners with multiple regions of contact along their length (Zhao *et al*., 2002 and Figure 1). Moreover, positive cooperativity is observed in the dephosphorylation of CheY-P by CheZ, indicating that the two proteins have structural inter-dependencies (Silversmith *et al*., 2008). The six suppressor mutations were restricted to two codons in CheY. The codons represent two amino acids located in the core CheY domain, a domain involved in the interaction with the CheZ core domain (see Figure 1). Based on the crystal structure of CheY bound to the phosphoryl group of BeF3 (10.2210/pdb1FQW/pdb), Met60 is in close proximity to Asp57, the site of phosphorylation. Thus, the substitution of a branched side chain at that location could potentially alter active site geometry and affect reaction kinetics. Ala42 is further away, but again substitution of a larger residue there could have effects that propagate to the active site.

Though not found directly at the surface of CheY, it is also possible that altered amino acids in the interior of CheY could influence key aspects of CheY core domain structure and interactions with CheZ at the CheY: CheZ interface. Based on this scenario, we propose a model with 3 features (see Figure 6):

1. The CheZ IDR is a dynamic moving structure with a characteristic structural motif, for example a rotating kink or propeller, that limits the effective volume occupied by the CheZ carboxyl terminus, affecting affinity for CheY and thus helping to “reel” in the CheY protein, B) the affinity of CheZ for CheY is limited when a key amino acid in the CheZ IDR (e.g., G188) is mutated, and C) the limited affinity of the IDR for CheY is partially or fully offset by mutations in the CheY core that facilitate improved interaction with CheZ. Since simple increased affinity between the core domains of CheZ and CheY could have been introduced through evolution through more straightforward mechanisms than introduction of a dynamic IDR reeling mechanism, we further propose that the IDR introduces advantages in chemotaxis signaling not apparent in a simple soft agar chemotaxis assay. Testing our suppressor mutants in more rigorous chemotaxis assays (quantitative chemotaxis assays) along with associated tethered cell assays to correlate chemotactic behavior with flagellar rotation, may disclose differences between wild-type and suppressors.

**Figure 6:**
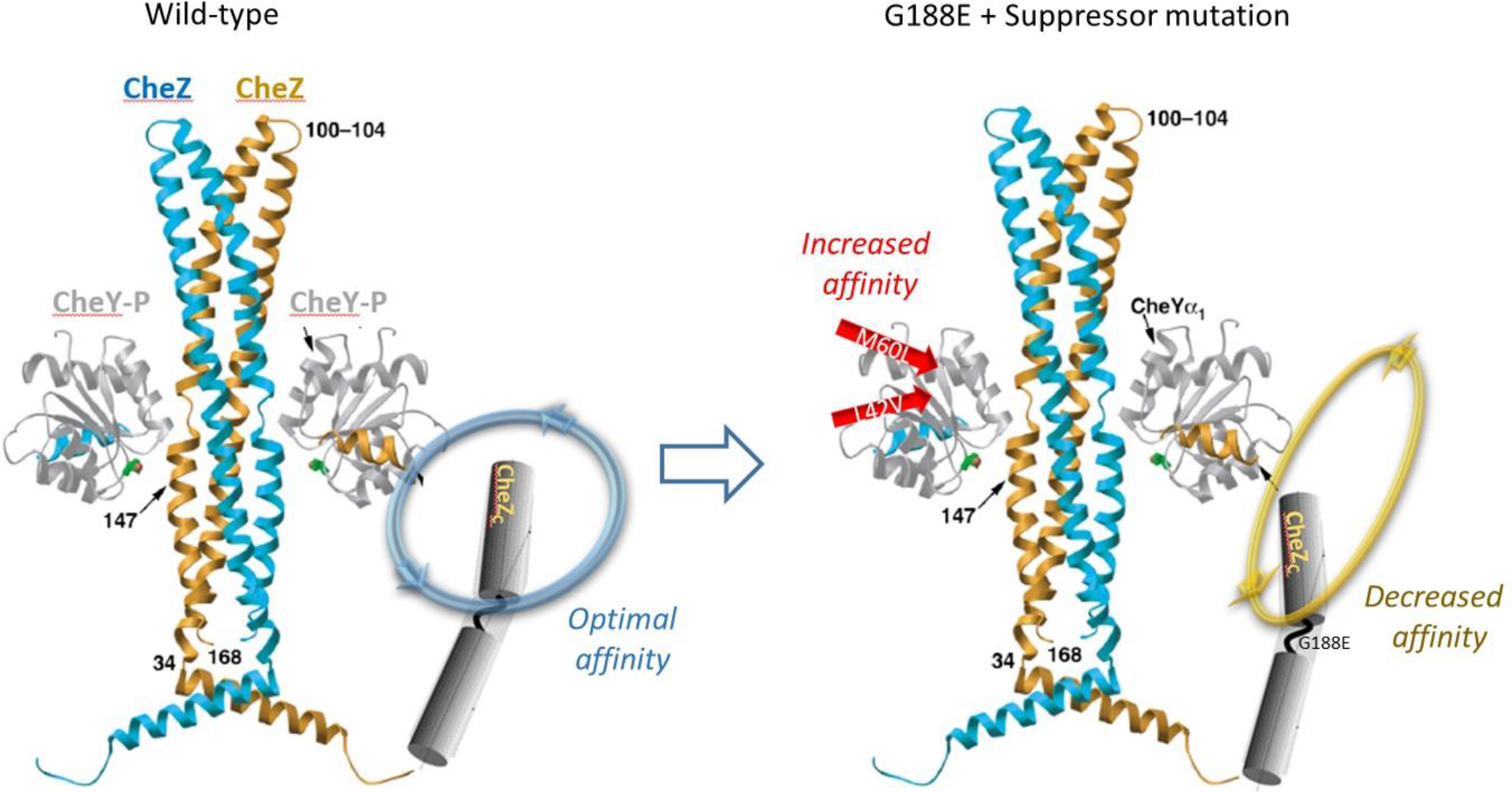
Model for restoration of activity of CheZ: CheY complex in presence of CheZ G188E containing a suppressor mutation in CheY, either at CheY M60 or L42 (see explanation in Discussion section). Blue arrow (wild-type) or yellow arrow (G188E plus suppressor) indicate a theoretical swivel/rotation of CheZ carboxyl terminus facilitated by the CheZ IDR. In this model, increased affinity of the CheZ and CheY cores containing suppressor mutations would compensate for decreased affinity resulting from mutated IDR.

In addition to further characterizing the chemotaxis phenotype of the suppressors, several more experiments are suggested by these studies. Biochemical properties of the CheY: CheZ phosphorylation reaction should be measured in the suppressor mutants versus wild-type proteins. It would be important to measure the detailed kinetics of phosphorylation of CheY since CheY-P has been shown to have a high “off rate” (Silversmith *et al*., 2008) that could be influenced by mutations in CheY. These studies also lay the foundation for future studies designed to test the generated hypotheses by assessing specific purified mutants in biophysical assays (e.g., stopped flow kinetics to directly measure rates of CheZ versus CheY-P binding). In sum, the genetic identification of CheY suppressor mutations provides the basis for testing specific hypotheses related to CheZ IDR structure: function.

## Acknowledgements

We acknowledge Dr. Verona Wagner (University of North Carolina, Chapel Hill, Department of Microbiology, Dr. Bourret lab) for contributing unpublished data on the CheZ IDR region which aided in the design of experiments described in this manuscript. We acknowledge Drs. Ruth E. Silversmith and Robert B. Bourret (University of North Carolina, Chapel Hill, Department of Microbiology and Immunology) for their guidance and expertise, access to relevant unpublished data, and for providing necessary *E. coli* strains.

